# The cerebellum does more than sensory-prediction-error-based learning in sensorimotor adaptation tasks

**DOI:** 10.1101/139337

**Authors:** Peter A. Butcher, Richard B. Ivry, Sheng-Han Kuo, David Rydz, John W. Krakauer, Jordan A. Taylor

## Abstract

Individuals with damage to the cerebellum perform poorly in sensorimotor adaptation paradigms. This deficit has been attributed to impairment in sensory-prediction-error-based updating of an internal forward model, a form of implicit learning. These individuals can, however, successfully counter a perturbation when instructed with an explicit aiming strategy. This successful use of an instructed aiming strategy presents a paradox: In adaptation tasks, why don’t individuals with cerebellar damage come up with an aiming solution on their own to compensate for their implicit learning deficit? To explore this question, we employed a variant of a visuomotor rotation task in which, prior to executing a movement on each trial, the participants verbally reported their intended aiming location. Compared to healthy controls, participants with spinocerebellar ataxia (SCA) displayed impairments in both implicit learning and aiming. This was observed when the visuomotor rotation was introduced abruptly (Exp. 1) or gradually (Exp. 2). This dual deficit does not appear to be related to the increased movement variance associated with ataxia: Healthy undergraduates showed little change in implicit learning or aiming when their movement feedback was artificially manipulated to produce similar levels of variability (Exp. 3). Taken together the results indicate that a consequence of cerebellar dysfunction is not only impaired sensory-prediction-error-based learning, but also a difficulty in developing and/or maintaining an aiming solution in response to a visuomotor perturbation. We suggest that this dual deficit can be explained by the cerebellum forming part of a network that learns and maintains action-outcome associations across trials.

**New and noteworthy:** Individuals with cerebellar pathology are impaired in sensorimotor adaptation. This deficit has been attributed to an impairment in error-based learning, specifically, from a deficit in using sensory prediction errors to update an internal model. Here, we show that these individuals also have difficulty in discovering an aiming solution to overcome their adaptation deficit, suggesting a new role for the cerebellum in sensorimotor adaptation tasks.

## Introduction

Visuomotor rotation tasks, which induce a discrepancy between movements of the limb and visual feedback, are a powerful tool for elucidating principles of motor learning (for a review see Krakauer, 2009). While learning in these tasks has typically been thought to be implicit, reflecting the training of an internal forward model via sensory prediction errors (Mazzoni & Krakauer 2006; Tseng et al., 2007; Schlerf et al., 2012), it has become clear that multiple processes can contribute to performance. These include reinforcement learning (Huang et al., 2011; Nikooyan and Ahmed, 2015; Galea et al., 2011), use-dependent plasticity (Diedrichsen et al., 2010; Verstynen and Sabes, 2011; Huang et al., 2011), and instructed aiming strategies (Mazzoni and Krakauer, 2006; Benson et al., 2011; Taylor and Ivry, 2011).

Most relevant to the current study is our recent work dissociating changes in performance that arise from modifications in aiming, which can be explicitly reported, and learning processes that are implicit, such as sensory-prediction-error-based learning (Taylor et al., 2014; Bond & Taylor, 2015; McDougle et al., 2015). The latter has been associated with the cerebellum, with compelling evidence coming from studies showing that individuals with cerebellar pathology display significant impairments in a range of sensorimotor adaptation tasks (Martin et al., 1996; Weiner et al., 1983; Rabe et al., 2009; Schlerf et al., 2013; Smith & Shadmehr, 2005). For example, in visuomotor rotation tasks, people with spinocerebellar ataxia (SCA) exhibit a reduced ability to counter experimentally-imposed perturbations. Importantly, they show attenuated aftereffects when the perturbation is removed (Schlerf et al., 2013; Werner et al., 2009). This is consistent with the idea that an intact cerebellum is required to compute sensory prediction errors via a forward model (Izawa et al., 2012; Miall & Wolpert, 1996; Haruno et al., 2001). This hypothesis is further supported by neuroimaging in humans (Schlerf et al., 2012) and neurophysiology in non-human species (Brooks et al., 2015) showing cerebellar activity that is correlated with sensory prediction errors.

In the context of a visuomotor rotation, we define aiming as choosing to move the hand towards a location other than the target, with the goal of having the cursor land on the target. *A priori*, there is no reason to believe that aiming is cerebellar dependent. Indeed, when individuals with cerebellar degeneration are provided with an explicit aiming strategy and visual cues to support the implementation of that strategy, they show near perfect performance in a visuomotor adaptation task (Taylor et al., 2010).

Given their ability to use an instructed strategy, it is puzzling that individuals with cerebellar damage are impaired in visuomotor rotation tasks when they are not provided with an explicit strategy. That is, why do they fail on their own to come up with an aiming solution to compensate for their impaired implicit learning, especially given that the perturbation results in a salient error that remains for many trials? This paradox suggests that the cerebellum may be necessary, not only for implicit learning, but also for discovering, implementing, and/or adjusting an appropriate aiming solution when it is not provided directly through instruction (see also, Vaca-Palomares et al., 2013).

We have developed a simple method to assess trial-by-trial fluctuations in aiming during visuomotor rotation tasks (Taylor et al., 2014) that allows for continuous assessment of aiming behavior and implicit learning. In the current set of experiments, we employed this method to determine the source(s) of impairment in individuals with cerebellar damage.

## Materials and Methods

### Participants

Experiment 1: Ten individuals with spinocerebellar ataxia (SCA, average age = 53.7, SD = 12.6 years; 2 female; 5 right-handed) were recruited for the study at the 2012 National Ataxia Foundation Annual Meeting (San Antonio, TX) and from the Berkeley, California community. SCA participants were only included if they did not exhibit clinical signs of cerebellar-type multiple system atrophy or evidence of moderate cognitive impairment. Four of the SCA group had a confirmed genetic subtype; for the others, the diagnosis was sporadic adult onset ataxia of unknown etiology (Table 1). The severity of ataxia was assessed with the International Cooperative Ataxia Rating Scale (ICARS, Trouillas et al., 1997). The SCA participants had an average ICARS score of 26.5 (SD = 14.7) out of a maximum score of 100. The participants were also screened for cognitive impairment with the Montreal Cognitive Assessment (MOCA; average = 28.1, SD = 1.5; all scored within normal range of 26-30) (Nasreddine et al., 2005).

**Table 1.** SCA neuropsychology and demographic information

| Participant | Gender | Age | Ed (years) | Handedness | Type | Diag (years) | Ataxia scale | MOCA |
| --- | --- | --- | --- | --- | --- | --- | --- | --- |
| **Abrupt** |  |  |  |  |  |  |  |  |
| ARSCA1 | Female | 63 | 16 | Left | SCA8 | 11 | ---- | 27 |
| ARSCA2 | Male | 73 | 20 | Left | SAOA | 4 | 20.5 | 28 |
| ARSCA3 | Male | 41 | 16 | left | SAOA | 19 | 27 | 29 |
| ARSCA4 | Female | 55 | 14 | Right | SCA2 | 13 | 48 | 26 |
| ARSCA5 | Male | 47 | 16 | Left | SAOA | 19 | 22 | 29 |
| ARSCA6 | Male | 42 | 16 | Right | SCA2 | 2 | 41 | 28 |
| ARSCA7 | Male | 62 | 19 | Right | SAOA/MSA | 4 | 12 | 30 |
| ARSCA8 | Male | 70 | 16 | Right | SAOA/Family Hist | 9 | 11 | 27.5 |
| ARSCA9 | Male | 44 | 22 | Left | SAOA/MSA | 2 | 12 | 30 |
| ARSCA10 | Male | 40 | 16 | Right | SCA2 | 11 | 45 | 26 |
| **Gradual** |  |  |  |  |  |  |  |  |
| GRSCA1 | Female | 35 | 17 | Right | SCA1 | 4 | 6* | 24 |
| GRSCA2 | Female | 30 | 17 | Left | SCA2 | 3 | 17* | ---- |
| GRSCA3 | Male | 44 | 17 | Right | SCA1 | 1 | 9* | 19 |
| GRSCA4 | Male | 79 | 13 | Right | SAOA | 6 | 11* | 14 |
| GRSCA5 | Male | 46 | 17 | Right | ADCA | 1 | 7* | 20 |
| GRSCA6 | Male | 41 | 17 | Right | SAOA | ---- | 23.5* | 23 |
| GRSCA7 | Male | 49 | 17 | Right | SAOA | 3 | 8* | 22 |
| GRSCA8 | Male | 36 | 17 | Right | SCA1 | 3 | 10.5* | 25 |
| GRSCA9 | Female | 69 | 19 | Left | SAOA | 8 | 3* | 30 |
| GRSCA10 | Female | 52 | 22 | Right | SCA6 | 14 | 1 | 28 |
| * SARA, otherwise ICARS |  |  |  |  |  |  |  |  |

Twelve age-matched control participants were recruited from the Berkeley or Princeton, New Jersey communities. These participants, based on self-reports, had no known neurological conditions. The data from two of the controls were not included in the final analysis: One consistently moved too slowly and the other failed to report the aiming locations on many trials (see below). Thus, the control group consisted of ten participants (average age = 59.7, SD = 14.7; SD = 1.4; 4 female; 10 right-handed). The control participants were also administered the MOCA (average = 27.6, SD = 2.2), with two scoring just below the normal range cutoff of 26 (23 and 25). They were included in the analyses given that their performance was similar to the other control participants on the experimental task.

Experiment 2: Twelve individuals with SCA were recruited from the Princeton community and from the Columbia University Medical Center (CUMC). These participants were selected after a clinical assessment determined that they did not exhibit symptoms of extracerebellar pathology (Parkinson’s disease, cerebellar-type multiple system atrophy). One individual was excluded from the analysis after failing to provide aiming reports, and a second was unable to complete the task in the time allotted, resulting in a final dataset of ten participants (average age = 48.1, SD = 16.2 years; average years of education = 17, SD = 1.6; 4 female; 8 right-handed; 6 confirmed genetic subtype; Table 1). The severity of ataxia symptoms was evaluated with the Scale for the Assessment and Rating of Ataxia Severity (Schmitz-Hubsch et al., 2006). The SCA participants had an average SARA score of 10.6 (SD = 6.2) out of a maximum score of 40. The SARA scale was used in experiment 2 reflected the preference of the CUMC neurologists. Due to time constraints, the MOCA was not administered to one of the SCA participants. Surprisingly, eight of the nine remaining SCA participants scored below the normal range on the MOCA (average score: 22.7, SD 4.8). In part, this may reflect the fact that, for four of these participants, English was a second language. It is also possible that this reflects a more severely compromised sample (although this is not supported by their ataxia scores), or more stringent scoring criteria. Ten age-matched control participants were recruited from the Princeton community (average age = 53.3, SD = 10.5 years; 5 female; 10 right-handed). The control participants were administered the MOCA (average = 25.5, SD = 1.1); five participants scored just below the normal range (three 25 and two 24).

Experiment 3: Twenty young adults were recruited from the research participation pool of the Department of Psychology at Princeton University (average age = 19.7, SD = 1.5; 9 female; 20 right-handed).

The experimental protocols were approved by the Institutional Review Boards at the University of California, Berkeley (experiment 1), CUMC (experiment 2), and Princeton University (experiments 2 and 3). All participants provided written informed consent. Participants in experiments 1 and 2 were paid an honorarium of $20 per hour, while participants in experiment 3 were compensated with class credit or $12.

### Experimental Apparatus

In all three experiments, participants made 7 cm horizontal reaching movements to visual targets. The targets were displayed on a 15-inch (Exp. 1) or 17-inch (Exps. 2 and 3) LCD monitor. The monitor was mounted horizontally, positioned approximately 25 cm above a digitizing tablet (Intuous Pro Large, Wacom). Given the position of the monitor, vision of the limb was occluded. All participants, regardless of handedness, were tested with their right hand. A digitizing pen was held in the right hand - regardless of handedness preference - and the movement required sliding the pen across the surface of the tablet. Feedback of hand position, when provided, was displayed in the form of a circular cursor displayed on the monitor.

### Procedure

We employed a variant of a visuomotor rotation task which requires participants to verbally report their aiming location on each trial. This procedure has been described in detail in a previous report (Taylor et al., 2014). At the start of each trial, a white ring was presented, indicating the distance of the participant’s hand from a start position (5 mm circle at center of screen). By continually making the ring smaller the participant could guide his or her hand to the start position. When the hand was within 1 cm of the center of the start position, the ring was replaced by a cursor, allowing the participant to precisely position the hand inside the start circle (Figure 1A).

**Figure 1.**
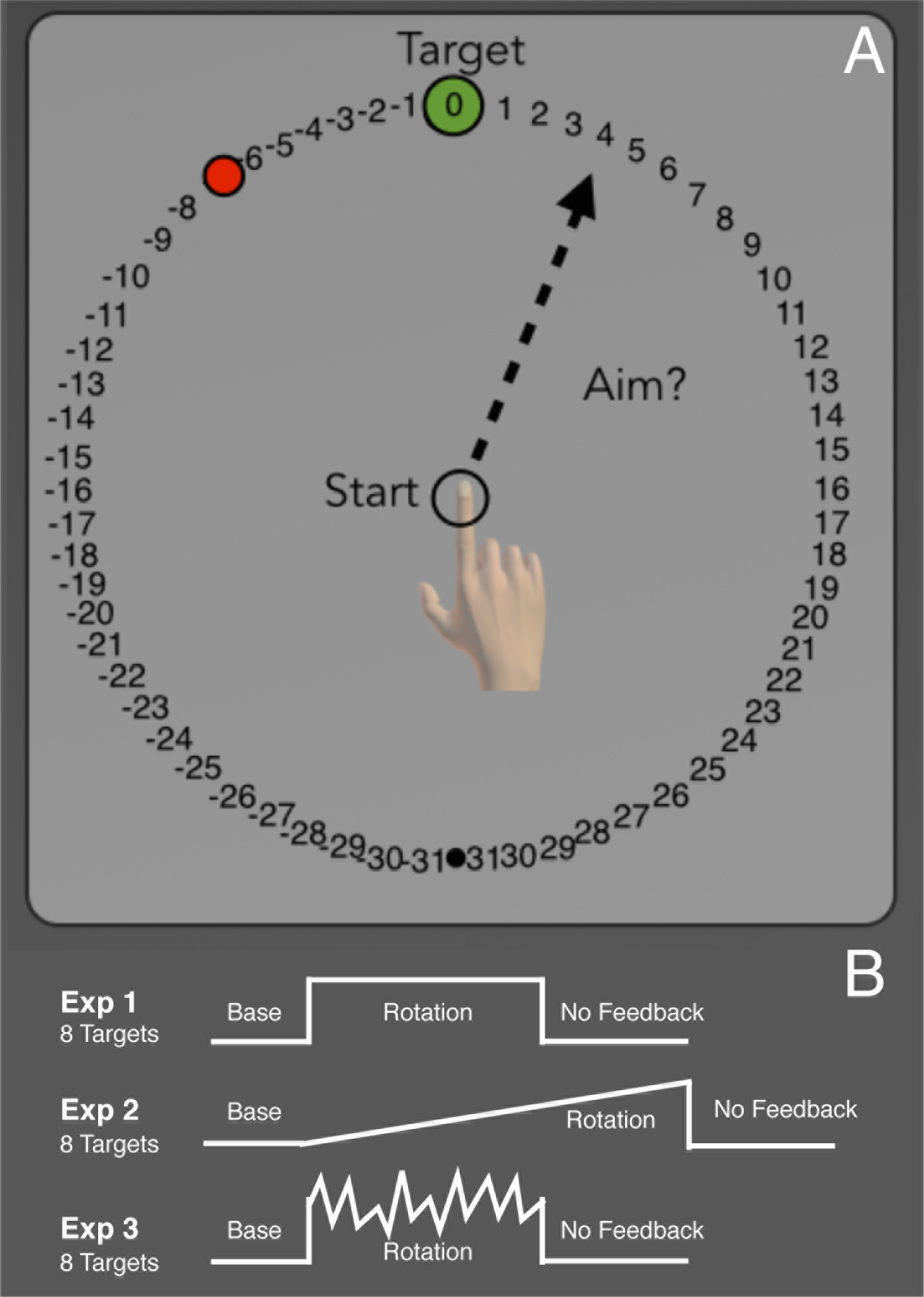
General task outline. A) On each trial a single target appeared at one of eight possible locations arranged equally around the start position. After the target appeared, but before moving, participants were asked to verbally report which number they were aiming towards for the current trial. B) In all three experiments, participants first completed 56 baseline trials with veridical feedback. This was followed by a rotation block in which the visual feedback was rotated about the start position. Finally, participants completed a no feedback washout block, where both visual and auditory feedback were withheld. In experiments 1 and 3 participants completed 128 rotation trials for a total of 264 trials, while in experiment 2 participants completed 220 rotation trials for a total of 456 trials.

After maintaining the start position for 1s, a green target circle (7 mm diameter) was presented. The target appeared at one of eight locations, separated by 45° along an invisible ring (radial distance from start circle of 7 cm). Each of the eight locations was presented within a block of eight trials and, within a block, the locations were randomly selected.

Participants were instructed to make a ballistic reaching movement, with the goal of getting the feedback cursor to appear at the target location. Participants were encouraged to reach beyond the target location, effectively shooting through the target. We chose to have the participants reach past the target to minimize the impact of intention tremor on accuracy, given that this symptom can become pronounced at the end of a rapid movement. Feedback was presented as an endpoint location in experiments 1 and 3. In these experiments, the cursor disappeared when the movement amplitude exceeded the 5 mm start circle and did not reappear until the amplitude reached 7 cm. Endpoint feedback was presented for 1.5 s at this location (subject to the visual perturbation— see below) in the form of a red cursor (3.5 mm diameter). In experiment 2, the cursor was visible during the outbound portion of the reach (online feedback). Once the amplitude reached 7 cm, the cursor position was frozen for 1.5 s. In all experiments, a pleasant “ding” sound was played if the feedback cursor intersected the target region; otherwise a mildly aversive “buzz” sound was played.

To encourage participants to make fast movements, a digital vocal sample saying “too slow” was played if the movement time was more than 400 ms. The movement protocol, involving an emphasis on fast movements with limited feedback (Exps. 1 and 3), was adopted to minimize feedback corrections, a problem for individuals with SCA (Tseng et al., 2007).

The visual workspace included a ring of numbered “landmarks”, spaced at regular intervals of 5.6° (Figure 1A). The numbers increased and decreased in the clockwise and counterclockwise directions, respectively, from the target. As such, the order of the landmarks rotated with the target. Prior to each movement on aiming report trials, the participant verbally reported the landmark they planned to reach towards. These verbal reports were recorded by the experimenter. Trials in which the participants failed to report their aim were excluded from the analysis. In experiment 1, the SCA participants failed to provide a report on 15.0% of the report trials, while control participants failed to provide a report on 1.8% of the trials. In experiment 2, the percentage of failed reports dropped to 1.6% and 0.4% for the SCA and control participants, respectively. In experiment 3, the college-age controls failed to provide reports on 1.1% and 1.0% of the trials for the No-Variance and Variance-Added groups, respectively. Refinements in the clarity of task instructions are likely responsible for the higher response rate in experiments 2 and 3.

The experiment was divided into five blocks: baseline, baseline-report, rotation, washout no-feedback, and washout with feedback. The participants first completed a baseline block of 48 trials with veridical feedback (Figure 1B). The report task was then described and participants completed eight trials, verbally reporting the aiming landmark prior to each reach. Feedback was veridical on these trials (and participants almost always reported the aiming location as “zero”).

The visual rotation was introduced in the rotation block. In experiment 1, this was a 45° counterclockwise rotation, imposed for 128 trials. In experiment 2, the rotation was introduced gradually over 320 trials, with 0.144° added on each trial until the full 45° counterclockwise rotation was achieved. For experiment 3, a counterclockwise rotation was present for 128 trials. For one group of participants (No-Variance), the size of the rotation was constant at 45°; for a second group of participants, the size of the rotation on a given trial was drawn from a Gaussian distribution with a mean of 45° and a standard deviation of 11° (Variance-Added). In all three experiments, participants reported the aiming location prior to making their reach.

Following the rotation block, participants made an additional 40 reaches, but the rotation, cursor feedback, and the aiming landmarks were removed (washout no-feedback). For this block, participants were explicitly instructed to aim directly to the green target. Neutral auditory feedback, in the form of a ‘knock’ sound, indicated when the reach amplitude exceeded 7 cm, but otherwise provided no information related to target accuracy. Veridical feedback was restored for a final 40-trial block (washout with feedback).

### Movement analysis

All initial data analyses were performed using Matlab (MathWorks) and statistical analyses were performed in SPSS (IBM, 2011). Task performance was assessed by calculating the angular difference between the target and the initial heading angle of the hand. A participants’ hand location could drift within the start circle during the report period of the trial, therefore we computed heading angle relative to the location where their hand left the start circle. This was done by fitting a straight line between samples taken at 1 and 3 cm from the start position (Taylor et al., 2014). We used initial heading angle rather than endpoint angle to allow for a visual comparison between the online and endpoint feedback conditions. The initial heading angle is also less susceptible to noise that might come about from the SCA participants’ ataxia.

For averaging across trials, movement trajectories were rotated to a common axis (e.g., as though the target was always located at 0°). With this convention, a positive angle indicates a deviation in the clockwise direction and a negative angle indicates a deviation in the counterclockwise direction. Note that the heading angles are reported in hand space rather than as target error. With this convention, hand heading angle will change in the opposite direction of the rotation as performance improves.

The mean hand heading angle was calculated on an individual basis for four different epochs: 1) The last eight trials of the baseline block, 2) the first and 3) last eight trials of the rotation block (early and late rotation), and 4) the first eight trials of the no-feedback washout block (washout). Trials were binned into groups of eight trials, we report the mean and standard error of the mean for each bin. Since the eight targets were presented in a random order within each cycle, this ensured that all targets were equally represented in each bin. In experiment 1, one participant with SCA failed to complete the last eight trials of the rotation block so the mean of their second to last bin of eight trials was used. We did not fit an exponential function to the time series of hand heading angles during the rotation block given the non-monotonic nature of the aim report data (Taylor et al., 2014).

To obtain an estimate of implicit learning, the reported aiming angle was subtracted from the measured hand heading angle on each trial. We refer to this measure as implicit learning, since it could contain a number of processes in addition to error-based updating of a forward model, such as use-dependent learning and reinforcement learning. The mean aiming angle and implicit learning was calculated for three different epochs: 1) The last eight trials of the baseline block, 2) the first and 3) last eight trials of the rotation block (early and late rotation).

In terms of kinematics, we measured velocity and movement time. Velocity was computed by submitting the hand position data to a fourth-order Savitzky-Golay filter (Savitzky & Golay, 1964; Smith et al., 2000). Movement onset could not always be based on the time at which the hand left the start circle because the hand occasionally wandered from this position during the report phase. As such, movement onset was estimated by a two-part procedure. We first identified the point in the time series where the movement amplitude reached 2 cm from the start position. From this point, the time series was searched backwards to find the time point when the participant either left the start circle, or when the movement started from outside the start position, the time point with the minimum radial distance from the start position.

We also excluded trials from the rotation block if the implicit learning estimate for that trial was more than 3 standard deviations outside the median implicit learning estimate for that participant. We used this criterion as a proxy to identify trials in which the participant may not have provided an accurate report of their aiming location, or that the movement itself was highly discrepant. Using this criterion, less than 1% of the trials were excluded for the SCA participants over experiments 1 and 2 and control participants in experiment 2. In experiment 1, control participants had 2.1% of trials excluded. For experiment 3, 1.5% and 1.6% of the trials were excluded for the Variance-Added and No-Variance groups, respectively.

### Power Analysis

We performed a power analysis to estimate the minimum sample size required to obtain an expected effect size, using the dataset from Taylor et al. (2010). Specifically, we focused on two “pure” measures of implicit learning, the extent of hand angle drift when participants were provided with an aiming strategy, and the magnitude of the aftereffect, comparing these measures between SCA and control participants. Power was estimated for an independent samples *t*-test, using a two-tailed *α* of 0.05 and power of 0.95. Based on the group means and standard deviations from the measure of implicit learning (i.e., drift) in Taylor et al. (2010), the effect size is *d* = 2.63 (*µ*_Control_ = 11.3°, *σ*_Control_ = 2.2°, *µ*_SCA_ = 5.9, *σ*_SCA_ = 1.9°). From this value, a minimum sample size of five participants is required in each group. A similar estimate of sample size is obtained using the aftereffect values (*µ*_Control_ = 6.2°, *σ*_Control_ = 2.4°, *µ*_SCA_ = 0.3, *σ*_SCA_ = 1.6°, *d* = 2.89). To be conservative, we doubled this number and recruited a minimum of 10 participants for each group for all of the experiments (Button et al., 2013).

## Results

### Experiment 1: Cerebellar damage impairs both aiming and adaptation to an abrupt rotation

Following a baseline block with veridical endpoint feedback, we introduced the aiming report task, requiring participants to indicate the number corresponding to the aiming location prior to each movement (Figure 1A). For the first eight trials in which feedback remained veridical (baseline-report block), participants in both groups tended to report aiming directly at the target (Control: −0.5 ± 0.3°; SCA: −0.3 ± 0.4°; group comparison: t_18_ = 0.5, p = 0.65). Consistent with their aiming reports, the heading angles of the reaches were directed towards the target with a small clockwise bias (Control: 2.9 ± 0.9°; SCA: 2.0 ± 1.8°; group comparison: t_18_ = 0.5, p = 0.65). In sum, the behavior during the baseline phase was similar between the two groups.

The introduction of the perturbation (rotation block) induced changes in heading angle from baseline for both groups (Figure 2A). To examine the initial phase of learning, we focused on the first eight trials. Participants with SCA displayed smaller heading angles (5.0 ± 3.8°) over these trials compared to the control participants (20.7 ± 3.5°; Figure 2D). The difference in performance was even more pronounced at the end of the rotation block. Over the final eight trials, the controls had almost completely countered the perturbation (41.3 ± 5.0°); in contrast, SCAs were only partially countering the perturbation (16.5 ± 7.9°). To determine whether there were any differences in performance over the course of the rotation block, we performed a mixed factorial repeated measures ANOVA with factors of Group (Control and SCA) and Time (Early Rot and Late Rot). A main effect of time is expected, since participants should compensate for more of the perturbation at the end of the rotation block than in the beginning. There are two, non-mutually exclusive, ways in which SCA participants’ performance could differ from controls. SCA participants might be generally impaired in compensating for the perturbation, relative to controls, in which case a main effect of group would be expected. Additionally, SCA participants could be slower to respond to the perturbation, in which case an interaction would be expected. The ANOVA revealed a main effect of group (F_(1,18)_ = 12.6, p = 0.002) and a main effect of time (F_(1,18)_ = 10.4, p = 0.005). No Group X Time interaction was present (F_(1,18)_ = 0.8, p = 0.37). Thus, the performance of both groups improved (cursor terminated closer to the target) over the course of the rotation block, and the performance improvement was considerably greater for the controls compared to the SCAs.

**Figure 2.**
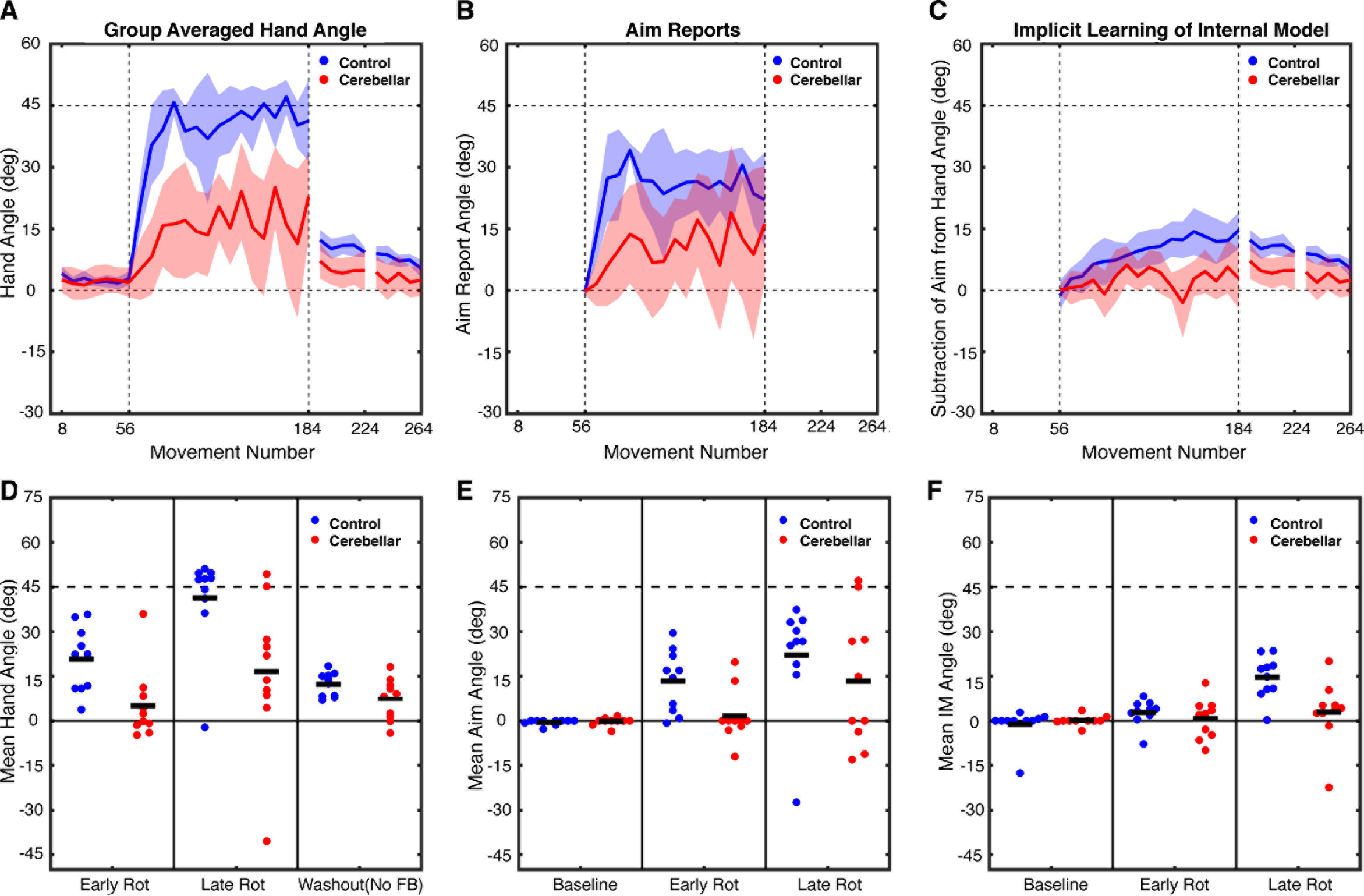
Experiment 1: Abrupt Rotation performance metrics. The top row depicts group averaged data for A) hand heading angle, B) aim reports, and C) estimates of implicit learning (Hand heading angle - Aim). The data are based on averages taken over bins of 8 trials for each participant, and then averaged across participants for each group. Shaded lines represent confidence intervals around the mean. Bins are marked with the trial number of the last trial of that bin. The bottom row shows individual participant data (dots) and group means (horizontal bar) for D) hand heading angle, E) aim report, and F) implicit learning estimate, with the data averaged over the first 8 trials of the rotation block, the last 8 trials of the rotation block, or the first 8 trials of no-feedback washout block.

To measure the size of the aftereffect, participants completed a no-feedback block in which they were instructed to aim directly to the target. Feedback in this block was limited to an auditory tone, indicating when the movement amplitude had exceeded 7 cm. A comparison of the first eight trials of this block to the 8 baseline trials revealed a reliable aftereffect for both controls (12.2 ± 1.3°; t_9_ = 5.5, p = 0.0004) and SCAs (7.1 ± 2.2°; t_9_ = 3.0, p = 0.02; Figures 2A, 2D). However, when comparing the two groups, the magnitude of the aftereffect was significantly larger in the control group compared to the SCA group (t_18_ = 2,0, p = 0.03). Additionally, our measure of implicit learning provides complementary evidence that this process is impaired in the SCA group (see below).

#### Verbal Reports

The time series of the aiming reports revealed that a large portion of the performance changes for the controls was due to a change in their reported aiming location (Figure 2B). For control participants, the mean aiming location was shifted from 13.3 ± 3.3° over the first eight trials to 22.1 ± 5.9° by the last eight trials of the rotation block. On average, these values were attenuated in the SCA participants (Figure 2E). Here, the early and late aiming angles were shifted from the target location by 1.6 ± 2.8° and 13.3 ± 7.0°, respectively. As with hand heading angle, to determine whether there were any differences in the verbal aiming reports over the rotation block, the aiming reports for early and late in the rotation block were submitted to a mixed factorial ANOVA with the same factors of Group (Control and SCA) and Time (Early Rot and Late Rot). The ANOVA revealed an effect of Time (F_(1,18)_ = 6.3, p = 0.03), suggesting aiming angles increased over the rotation block, but only a trend for an effect of Group (F_(1,18)_ = 3.2, p = 0.09). No Group X Time interaction (F_(1,18)_ = 0.1, p = 0.74) was present. From visual inspection of the aiming time series, control participants appeared to have an initial large shift in aiming angle, which then began to decrease slowly over the course of the rotation block. Given this non-monotonic nature of the aiming report time series, as well as the high variance in the initial trials, we performed an additional *post-hoc* analysis comparing the aim reports for the two groups across the whole rotation phase. On this composite measure, the aiming reports indicated larger shifts in aiming location for control (25.7 ± 3.8°) compared to SCA participants (11.2 ± 5.1°; t_18_ = 2.3, p = 0.035).

To estimate implicit learning, we subtracted the reported aiming angle from the hand heading angle for each trial (Figure 2C). We again focused on the mean for the first and last eight trials. For control participants, the estimate of implicit learning increased from 2.8 ± 1.4° early in the rotation block to 14.6 ± 2.3° by the end. By comparison, SCA participants had an estimate of no implicit learning (0.6 ± 2.1) early in the rotation block, which only barely increased to 2.9 ± 3.4° by the end of the block. As was done for hand heading angle and the verbal aim reports, to compare changes in implicit learning over the rotation block, these values were submitted to an ANOVA with factors of Group and Time. The ANOVA revealed significant main effects of Group (F_(1,18)_ = 5.6, p = 0.03), resulting from generally impaired implicit learning for SCA participants, and Time (F_(1,18)_ = 17.7, p = 0.001), due to implicit learning estimates increasing from early to late in the rotation block. A Group x Time interaction (F_(1,18)_ = 8.1, p = 0.01) was also present, suggesting the impaired implicit learning in SCA participants differed relative to control participants between early and late in the rotation. As can be seen in Figure 2F, the estimate of implicit learning was markedly lower for the SCAs, an effect that was especially pronounced late in the rotation block. Thus, using both the estimate of implicit learning during the rotation block and the aftereffect measure, adaptation was impaired in SCA participants compared to controls.

In summary, the individuals with spinocerebellar ataxia exhibited a performance impairment when presented with a 45° visuomotor rotation, similar to that observed in previous studies of sensorimotor adaptation (e.g., Martin et al., 1996; Schlerf et al., 2013). By obtaining verbal reports, we were able to dissociate adjustments in aiming from implicit learning. The results indicate a dual-impairment: Not only was implicit learning attenuated in the SCA participants, but they also tended to aim to locations closer to the target during the early stages of the rotation block compared to controls. While SCA participants did modify their aim, these adjustments failed to effectively counter the rotation such that their overall performance only compensated for about half of the perturbation by the end of training. Thus SCA participants were impaired in self-discovery of an aiming strategy, which stands in contrast to our previous finding that SCA participants are quite competent in carrying out strategy if it is provided by instruction (Taylor et al 2010).

### Experiment 2: Gradually introducing a rotation fails to alleviate the impairment in aiming and adaptation as a result of cerebellar damage

With an abruptly introduced rotation, there appears to be two stages of aiming: an initial large change in aim to compensate for the salient error induced by the perturbation, followed by small trial-to-trial adjustments to maintain accurate performance in the presence of small errors. It is likely that this first initial stage is the selection and implementation of an explicit general aiming strategy, however, the extent to which this second smaller adjustment phase is explicitly generated is less clear. It may be that the small trial-to-trial adjustments are achieved by something like implicit aiming, where an implicit mechanism is used to generate the adjustment to which explicit access is gained afterwards. Providing an instructed aiming strategy would direct the initial large change in aim, however, as adaptation increases and compensates for more of the perturbation, to maintain accurate performance and offset adaptation, small adjustments in aim may be necessary from trial-to-trial. Given that when SCA individuals are provided with an instructed aiming strategy they can counter an abrupt rotation, and maintain performance, they may not be impaired when only small trial-to-trial changes in aim are necessary. To investigate this, in Exp. 2, we introduced the perturbation gradually over the rotation block. Note, we extended the rotation block from 128 to 320 trials and provided online cursor feedback as the participants reached towards the target to increase the contribution of implicit learning.

For the first eight trials of aiming under veridical feedback (baseline-report block), participants in both groups tended to report aiming towards the target (Control: −2.4 ± 2.2°; SCA: −0.9 ± 1.2°; group comparison: t_18_ = 0.6, p = 0.56). Consistent with their aim reports, the heading angles for these first eight baseline trials were directed towards the target with a small clockwise bias (Control: 1.4 ± 0.6°; SCA: 3.9 ± 1.6°; group comparison: t_18_ = 1.5, p = 0.15). As in experiment 1, the emphasis on participants making quick slicing movements towards the target resulted in relatively similar behavior for both groups of participants.

Following the baseline block, a 45° counterclockwise visuomotor rotation was gradually introduced over 320 trials. The small perturbation (only 1.15° on the 8th trial) did not induce reliable changes from baseline in hand heading angle over the first eight trials of the rotation block for either the controls (−0.7 ± 2.8°) or the SCA participants (1.6 ± 1.2°; Figure 3D). By the end of the rotation block, when the full 45° rotation was present, both groups had adjusted their hand heading angles in response to the perturbation (Figure 3A). This change, averaged over the last eight trials, was substantially larger in the controls (35.8 ± 1.3°) compared to the SCA participants (18.2 ± 4.2°). As in experiment 1, to compare performance over the rotation block these values were submitted to a mixed factorial ANOVA. The ANOVA revealed a main effect of Group F_(1,18)_ = 8.8, p = 0.008), a main effect of Time (F_(1,18)_ = 93.9, p < 0.0001) and a Time X Group interaction (F_(1,18)_ = 13.3, p = 0.002). Thus, as in experiment 1, the main effect of Group reveals that SCA participants compensated less for the perturbation than did control participants, showing that their performance impairment is observed with both abrupt and gradual perturbations (see Schlerf et al., 2013). Additionally, the presence of an interaction suggests that the relative impairment of SCA participants to controls differed between early and late in the rotation. Visual inspection suggests the difference in performance was larger late in the rotation block, which is expected given that the gradual introduction of the perturbation resulted in only a ~1° perturbation during this early phase.

**Figure 3.**
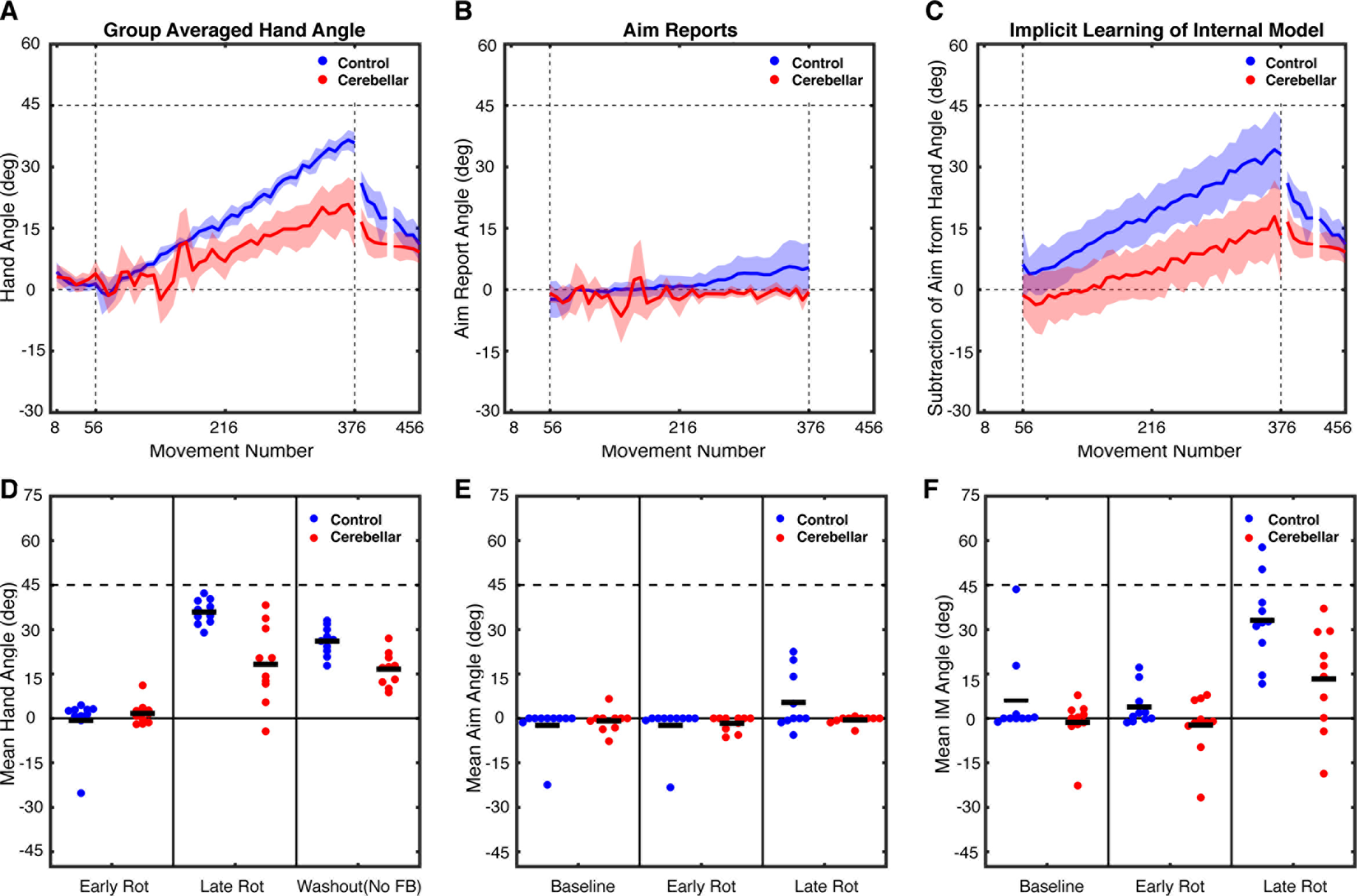
Experiment 2: Gradual Rotation performance metrics. The top row depicts group averaged data for A) hand heading angle, B) aim reports, and C) estimates of implicit learning (Hand heading angle - Aim). The data are based on averages taken over bins of 8 trials for each participant, and then averaged across participants for each group. Shaded lines represent confidence intervals around the mean. Bins are marked with the trial number of the last trial of that bin. The bottom row shows individual participant data (dots) and group means (horizontal bar) for D) hand heading angle, E) aim report, and F) implicit learning estimate, with the data averaged over the first 8 trials of the rotation block, the last 8 trials of the rotation block, or the first 8 trials of no-feedback washout block.

On the no-feedback washout block, cursor feedback was withheld and the participants were instructed to aim directly for the target. Comparing the first eight trials of this block to the eight baseline trials revealed a reliable aftereffect for the controls (26.0 ± 1.5°; t_9_ = 15.8, p < 0.0001) and SCAs (16.6 ± 1.8°; t_9_ = 5.2, p = 0.0005). However, the magnitude of the aftereffect was smaller in SCA compared to control participants (t_18_ = 4.0, p = 0.0004; Figures 3A, 3D). Thus, on this measure of implicit learning, the SCA participants were impaired relative to the control group.

#### Verbal Reports

In contrast to experiment 1, the time series of the aiming reports revealed that only a small portion of the change in hand heading angle for the controls was due to a change in their aiming location (Figure 3B). Over the first eight trials of the rotation block, their mean aiming location was −2.4 ± 2.3°. Note that this shift is in the counterclockwise direction and would effectively increase the perturbation; we assume this reflects noise or an attempt to negate the effects of intrinsic bias (Gibo et al., 2013; Vindras et al., 1998 & 2005; Ghilardi et al., 1995). By the last eight trials, the aim was shifted by 5.3 ± 3.1° in the clockwise direction, effectively helping to counter the rotation. In contrast, the SCA participants did not consistently shift their aiming location over the course of the rotation block (Figure 3E). Compared to baseline, they showed a shift of −1.7 ± 0.8° over the first eight trials and a shift of −0.6 ± 0.4° over the last eight trials. When comparing the verbal aiming reports with an ANOVA, the effect of Time was significant (F_(1,18)_ = 6.1, p = 0.024), but the effect of Group was not (F_(1,18)_ = 1.5, p = 0.24). There was a trend towards a Time X Group interaction (F_(1,18)_ = 3.3, p = 0.085).

We performed two additional post-hoc comparisons to quantify the extent of aiming in the control and SCA participants. First, to determine whether aiming direction changed over the course of the rotation block, a within subject *t*-test was conducted, comparing aiming during the last 8 baseline trials and the last 8 rotation trials. By this measure, the control participants adjusted their aim by the end of the rotation block (t_9_ = 2.2, p = 0.051), although this was only marginally significant. In contrast, the SCA participants did not exhibit a reliable shift in aim (t_9_ = 0.3, p = 0.79). Second, to compare overall aiming between the two groups, a *t*-test was performed comparing the aim reports averaged across the entire rotation phase. Using this measure, we observed a reduced shift in aiming direction for the SCA group compared to the controls (t_18_ = −2.1, p = 0.047). Thus, despite the relatively small changes in aiming observed for the control participants, these comparisons suggest larger changes in aim for the control participants compared to SCA participants. Indeed, the SCA participants failed to adjust their aim in a consistent manner during the rotation block, despite the fact that the target error became quite pronounced by the end of the block.

We employed the subtractive procedure to estimate implicit learning (Figure 3C), and focused on the first and last eight trials during the rotation block for our statistical analysis (Figure 3F). For control participants, the estimate of implicit learning increased from 3.8 ± 2.1° early in the rotation block to 33.1 ± 4.5° by the end. By comparison, SCA participants had an estimate of no implicit learning (−2.3 ± 3.1) early in the rotation block, which increased to 13.2 ± 5.5° by the end of the block. An ANOVA on these values revealed main effects of both Group (F_(1,18)_ = 7.0, p = 0.016) and Time (F_(1,18)_ = 59.7, p < 0.0001). The main effect of group results from smaller implicit learning estimates overall for SCA participants compared to controls. In addition, the Group X Time interaction was significant (F_(1,18)_ = 5.6, p = 0.030), reflecting the fact that the control participants had a larger increase in implicit learning over the course of the rotation block compared to the SCA participants. This result converged with that observed in the measure of the aftereffect, replicating the impaired adaptation for SCA participants shown in experiment 1.

While our results are comparable across experiments 1 and 2, we caution against drawing any strong inferences from any differences in results between the experiments. First, there is the problematic nature of null results (the lack of a difference between controls and SCA participants on the aftereffect data in experiment 1). Second, there are substantial differences between the two tasks. In experiment 2, participants completed more than twice as many rotation trials as in experiment 1, and received online cursor feedback. We would expect both factors to enhance implicit learning (in controls), offering greater sensitivity when comparing their performance to the SCA participants.

The results of experiment 2 demonstrate that the SCA participants were impaired in responding to a 45° gradual visuomotor rotation. As in experiment 1, their deficit appears to be manifest in measures of both implicit learning and aiming. The aiming deficit was apparent in experiment 2, even though only small adjustments in aiming location are necessary from trial-to-trial to maintain performance. We note that, for both groups, the amount of aiming was markedly attenuated in this experiment, and this may have contributed to the fact that the target error remained substantial at the end of the experiment. For controls, the final error was around 9°; for the SCA group, the final error was around 27°. Despite this large error, the SCA participants failed to alter their aiming locations; they were unable to compensate for their impairment in implicit learning by deploying an aiming solution.

Given the variable performance of SCA participants in experiments 1 and 2, and that SCA leads to highly heterogeneous damage to the cerebellum, it may be tempting to map behavior to damage in cerebellar subregions or with specific subtypes of SCA. However, given this variability, a larger sample size than we have here would be necessary (see Kansal et al., 2016). With the current dataset, any conclusions involving SCA subtypes, or more specific regions of the cerebellum, would likely be driven by lesions and performance in only a few participants, where lesion reconstruction is unlikely to yield reliable results (Rorden et al., 2007; Kimberg et al., 2007). Additionally, correlating behavioral deficits to damage in cerebellar subregions is likely to be best tested in individuals with focal lesions, where the pathology is more localized than the broad deterioration in SCA.

### Experiment 3: Higher motor variability does not account for the aiming and adaptation deficits due to cerebellar damage

A feature of spinocerebellar ataxia is the presence of increased movement variability. Indeed, even though we focused on the initial heading angle in experiments 1 and 2, the reaching movements for the SCA participants were more variable than the controls. For example, limiting the analysis to the initial baseline block of experiment 1 (before the aiming task was introduced), the standard deviation of the heading angles for the SCA and control groups were 11° and 5°, respectively. This increase in movement variability may make it difficult for individuals with SCA to develop a reliable aiming solution because they cannot converge on a consistent direction. To make this concrete, consider the situation where an individual, after encountering a 45° clockwise perturbation as in experiment 1, opts to aim to a location that is 30° in the counterclockwise direction. Evaluating the utility of this aiming solution will be hampered if the reach deviates widely from the selected trajectory (independent of adjustments induced by implicit learning). Similarly, increased motor variability in experiment 2 might make it difficult for the SCA participants to gauge the effects of the increasing perturbation.

The impact of motor variance on learning has been explored in previous studies of visuomotor adaptation (Wu et al., 2014; Therrien et al., 2015, He et al., 2016). In terms of the effects of cerebellar pathology, Schlerf et al. (2013) found that individuals with SCA exhibited impaired implicit learning, even when one considers how increased motor variability might impinge upon learning and performance. Here we ask how an increase in motor variability might influence the discovery of an aiming solution. Rather than create conditions in which we directly manipulate motor variability, we added noise to the movement feedback presented to adults and examined the effect this had as they learned to respond to a 45° perturbation.

College-aged adults were randomly assigned to one of two groups in experiment 3. In the No-Variance group, the procedure was identical to experiment 1 with the participants exposed to a constant 45° counterclockwise perturbation during the rotation block. In the Variance-Added group, we (crudely) simulated the effects of ataxia by pseudo-randomly varying the size of the rotation on each trial during the rotation block (Figure 4). To this end, a noise term was added to the 45° perturbation. The size of the rotation on each trial was based on a random sample from a Gaussian distribution with a mean of 45° and standard deviation of 11°. The value of 11° was chosen because it is the mean of the individual standard deviations of the hand heading angle for the SCA participants in Experiment 1. Given that feedback is limited to the reach endpoint (at 7 cm), the participants in the Variance-Added group would experience a noisy 45° perturbation.

**Figure 4.**
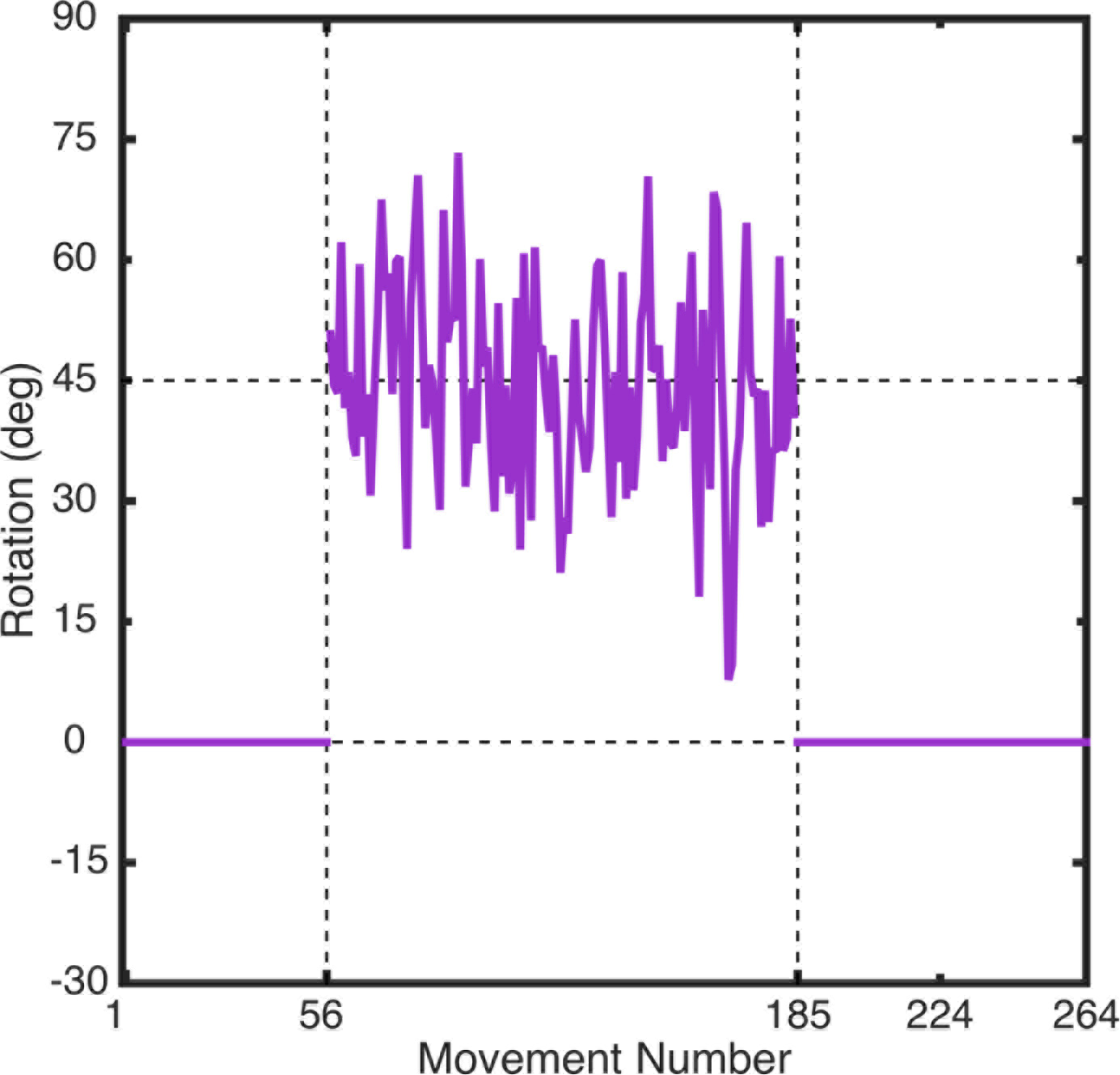
Sample perturbation schedule for Variance-Added group in experiment 3. The visuomotor rotation for each trial in the rotation block was was drawn from a gaussian distribution with a mean of 45° and standard deviation of 11°. The variance of the gaussian was based on the mean variance in hand heading angle seen in participants with ataxia in experiment 1. Participants in the No-Variance control group experienced a constant 45° rotation on all trials in the rotation block (not shown).

During the initial baseline block (prior to reporting aim) the two groups had similar standard deviations of their heading angles (No-Variance: 6.3 ± 0.4°; Variance-Added: 6.4 ± 0.9°; t*18* = 0.04, p = 0.97), suggesting there were no baseline differences in movement variance between the two groups. For the first eight trials of aiming with veridical feedback (baseline-report block), the participants in both groups tended to report aiming towards the target (No-Variance: −1.1 ± 0.6°; Variance-Added: 0.1 ± 0.2°; group comparison: t_18_ = 0.6, p = 0.56). The hand heading angles were in the direction of the target, although the Variance-Added group exhibited a significant, albeit small clockwise bias (2.7 ± 0.7°) that was not observed in the No-Variance group (0.0 ± 0.5°, t_18_ = 3.1, p = 0.01). Given that the two groups received identical (veridical) feedback in these first two blocks, this difference likely represents chance variation in the participants’ baseline reaching bias (Ghilardi et al., 1995).

The 45° perturbation was introduced in the rotation block, along with the increase in endpoint variance for the Variance-Added group. Both groups displayed adjustments in heading angle in response to the perturbation (Figure 5A). These adjustments were similar for the No-Variance and Variance-Added participants, both early (No-Variance mean: 12.6 ± 4.5°; Variance mean: 11.7± 6.2°) and late in the rotation block (No-Variance mean: 44.2 ± 1.2°; Variance-Added mean: 41.8 ± 6.6°; Figure 5D). As in previous experiments, a mixed factorial ANOVA was used to compare learning over the rotation blocks, although in this case the factor of Group is comparing Variance-Added and No-Variance groups. There was a main effect of Time (F_(1,18)_ = 76.1, p < 0.0001), but no effect of Group (F_(1,18)_ = 0.07, p = 0.80) nor a Group X Time interaction (F_(1,18)_ = 0.04, p = 0.84), thus no differences in performance were present between the groups. The groups also exhibited similar aftereffects in the no-feedback block in which they were instructed to aim directly for the target. Both groups exhibited a significant aftereffect (No-Variance: 6.3 ± 0.9°; t_9_ = 4.8, p = 0.0009; Variance-Added: 7.9 ± 0.8°; t_9_ = 5.2, p = 0.0006), and the magnitude of the aftereffect was similar for the two groups (t_18_ = 1.4, p = 0.90). These results suggest that the imposition of added endpoint variance did not impact overall performance in response to a visuomotor perturbation.

**Figure 5.**
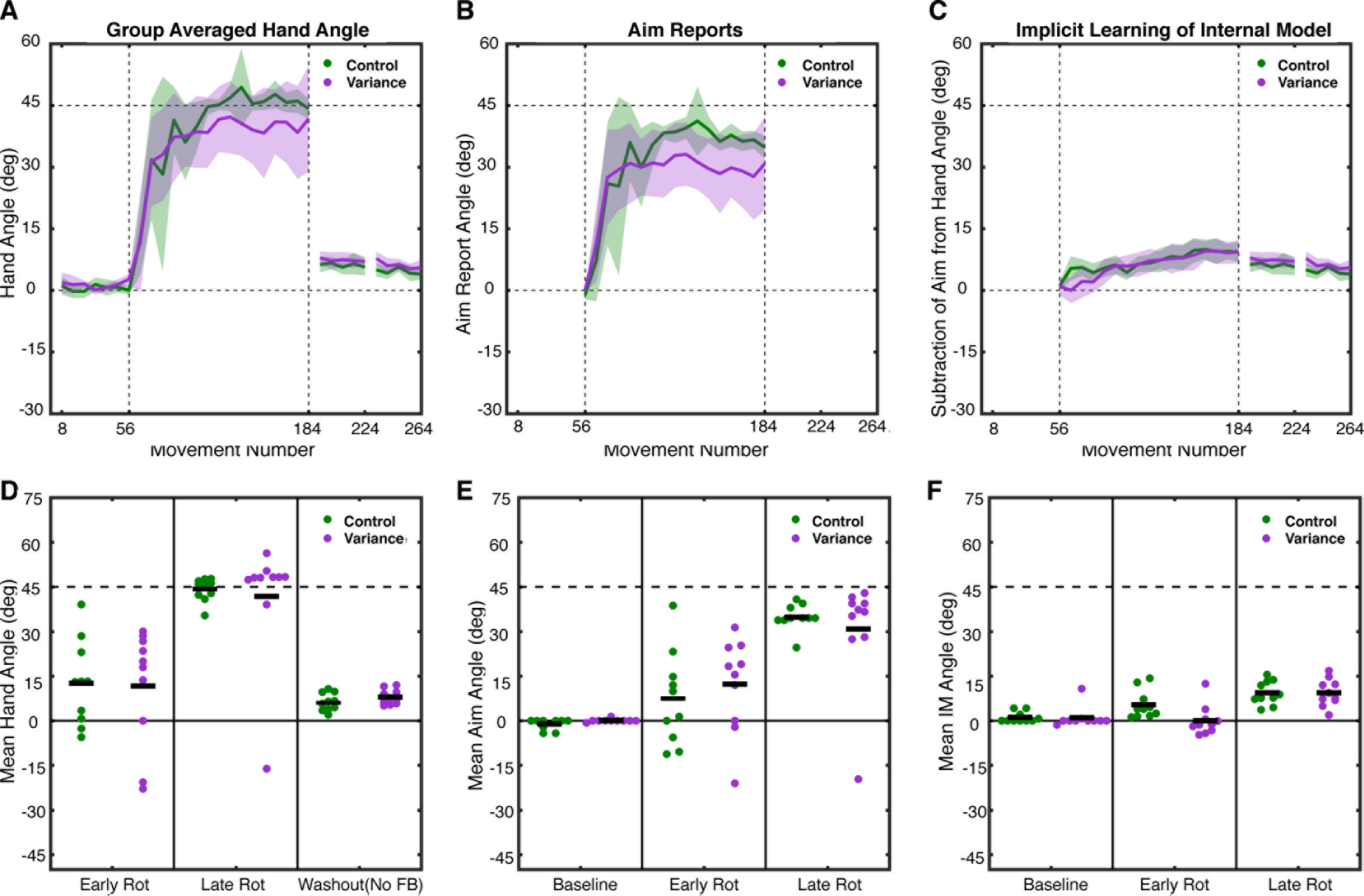
Experiment 3: Abrupt Rotation with variable rotation performance metrics. The top row depicts group averaged data for A) hand heading angle, B) aim reports, and C) estimates of implicit learning (Hand heading angle - Aim). The data are based on averages taken over bins of 8 trials for each participant, and then averaged across participants for each group. Shaded lines represent confidence intervals around the mean. Bins are marked with the trial number of the last trial of that bin. The bottom row shows individual participant data (dots) and group means (horizontal bar) for D) hand heading angle, E) aim report, and F) implicit learning estimate, with the data averaged over the first 8 trials of the rotation block, the last 8 trials of the rotation block, or the first 8 trials of no-feedback washout block.

Per experimenter instructions, participants in both groups moved rapidly, and there was no difference in movement duration between the groups during the rotation block (No-Variance: 274 ± 57 ms; Variance-Added: 214 ± 14 ms; t_18_ = 1.02, p = 0.32).

#### Verbal Reports

The aiming report data reveals that a large portion of learning was due to an adjustment in aiming location (Figure 5B), and that this effect was comparable for the two groups. For the No-Variance group, the mean aiming location was shifted 7.3 ± 5.0° over the first eight trials, and increased to 34.8 ± 1.4° by the last eight trials of the rotation block. Participants in the Variance-Added group had an initial shift of 12.3 ± 5.0°, which increased to 30.8 ± 5.8° by the last eight trials of the rotation block. A mixed factorial ANOVA comparing aiming over the rotation block revealed the main effect of Time was significant (F_(1,18)_ = 33.3, p < 0.0001), with participants increasing the angle of their aiming over the course of the rotation block (Figure 5E). However, there was no main effect of Group (F_(1,18)_ = 0.01, p = 0.92) nor a Group X Time interaction (F_(1,18)_ = 1.3, p = 0.27). While the overall pattern suggests that the aiming shift over the entire rotation block was larger for the No-Variance group (33.7 ± 2.3°) compared to the Variance-Added (29.0 ± 3.9°) group, this difference was not significant (comparison of means taken over the whole rotation block: t_18_ = 1.0, p = 0.31). Indeed, the group difference reflects the performance of one participant in the Variance-Added group who only reported small aiming angles.

Implicit learning was estimated for each trial by subtracting the aiming angle from the hand heading angle (Figures 5C, 5F). Participants in the No-Variance group had an estimate of implicit learning that increased from 5.3 ± 1.5° early in the rotation block to 9.4 ± 1.3° by the end. In the Variance-Added group participants had an estimate implicit learning that increased from 0.0 ± 1.6 early in the rotation block to 9.4 ± 1.4° by the end of the block. Implicit learning was compared with an ANOVA based on the first and last eight trials of the rotation block, revealing a main effect of Time (F_(1,18)_ = 26.0, p = 0.0001), but no main effect of Group (F_(1,18)_ = 2.9, p = 0.11) and only a trend for an interaction between these factors (F_(1,18)_ = 4.2, p = 0.055). While the group effect and interaction were not significant, a post-hoc comparison of the groups early (t_18_ = 2.5, p = 0.02) and late (t_18_ = 0.0, p = 0.99) in the rotation block, suggests that while participants in the Variance-Added group may have initially had less implicit learning early during the rotation block, their implicit learning caught up to the No-Variance group by the end of the rotation. When a further post-hoc comparison is made of implicit learning estimates averaged across the entire rotation block, no difference was present between the No-Variance group (7.2 ± 0.9) and the Variance-Added group (6.5 ± 1.2; t_18_ = 0.5, p = 0.62), confirming no difference in overall implicit learning was present.

In summary, the results of experiment 3 show that the injection of random endpoint noise had only a modest effect on performance. Interestingly, the effect of noise was, at least in terms of overall performance, limited to a possible slight reduction in the magnitude of implicit learning as estimated early during the rotation block, but no differences were present late in the rotation block, nor in aftereffect. There were also differences between the groups in how they modified their aim on a trial-by-trial basis, with the Added-Variance group continually modifying their aiming location in response to the added noise. Nonetheless, these participants were able to learn the mean of the 45° rotation and they changed their mean aiming angle to a similar degree as participants who experienced a constant perturbation. Taken together, these results suggest that an increase in endpoint variability does not result in profound deficits in implicit and aiming components of visuomotor learning. Thus, the difficulty exhibited by individuals with SCA in adopting an appropriate aiming solution is likely unrelated to their increased movement variability.

## Discussion

Previous work has repeatedly demonstrated that individuals with damage to the cerebellum are impaired in sensorimotor learning. The emphasis in this literature has focused on implicit deficits in error-based learning. However, here we have shown that SCA also leads to an impairment in the ability to discover and maintain an aiming solution to offset a visuomotor perturbation.

### Impaired aiming in SCA

To study the use of aiming behavior, participants reported their aiming location prior to each reach (Taylor et al., 2014). This task provides a direct probe on the use and evolution of an aiming solution and, by subtractive logic, a continuous estimate of implicit learning. We observed a dual-deficit in participants with SCA when presented with a visuomotor rotation. Not only did the SCA group exhibit impaired implicit learning, they also showed a failure to aim to counter the observed target error. They adjusted their aim across the perturbation block, choosing locations that tended to be in the appropriate direction to counter the rotation, but failed to fully compensate for the perturbation.

This aiming deficit helps answer the question posed in the Introduction: If people can compensate for a rotation through a multiplicity of processes, why do individuals with SCA not compensate for an implicit learning deficit through an increased reliance on aiming? The current results indicate that the ability to discover an aiming solution is also compromised.

This dual-deficit was not only observed when a 45° perturbation was introduced abruptly (Exp 1), but also when the 45° perturbation was introduced gradually (Exp 2). It has been assumed that aiming makes a minimal contribution to performance when the perturbation is introduced in a gradual manner. However, this assumes that implicit learning – the form arising from sensory-prediction-error-based learning – will be sufficient to compensate for the perturbation. The current data, as well as a recent report (Bond and Taylor, 2015), suggest that implicit learning is insufficient to achieve good performance when the perturbation is large (for controls, as well as SCA participants), and thus, the endpoint error will grow over the course of the rotation block. In experiment 2, the control participants began to adjust their aim, with deviations in the aiming location becoming consistent after approximately 150 trials into the rotation block. In contrast, the SCA group failed to invoke compensatory aiming even when the error became quite large. A priori, we might have expected the SCA participants to have larger aiming angles than the controls in order to compensate for their impaired implicit learning.

Taken together, experiments 1 and 2 demonstrate that SCA participants have difficulty adjusting their aim to counter a visuomotor rotation. Thus, in addition to an impairment in sensory-prediction-error-based learning, these individuals have difficulty using the feedback to develop and implement an appropriate aiming solution. Below, we consider possible explanations for this impairment in aiming.

### Increased movement variability does not account for impairments in aiming

In experiment 3, the addition of trial-by-trial noise to a constant perturbation was used to test the hypothesis that the observed aiming impairment is due to ataxia-related movement variability. This manipulation, however, only had a modest effect on the performance of the young healthy adults. Most relevant to the current study, the additional variability did not produce a significant effect on either overall performance or on aiming behavior. The Variance-Added group altered their aiming direction shortly after the onset of the (noisy) perturbation. While they continued to make aiming adjustments across the rotation block, the asymptotic size of the aiming shift was statistically indistinguishable from that observed in the No-Variance group.

We recognize that our noise manipulation is, at best, a weak approximation of the consequences of ataxia. We assume the college students attribute the increased variability to the environment, and not their motor system. Moreover, the increased variance is transient, unlike the chronic nature of ataxia. Nonetheless, the results of experiment 3 suggest that an inability to develop an appropriate aiming solution is not solely due to the motor variability associated with ataxia.

The Variance-Added group did exhibit a small, reduction in the estimate of implicit learning early the rotation block. However, there was no difference in the implicit learning estimate by the end of the rotation block, nor when compared across the entire rotation block. However, this delayed onset of implicit learning is likely distinct from the implicit learning deficit observed in SCA. Under conditions of high external noise, the learning rate should be reduced (Kalman, 1960; He et al., 2016). Moreover, using a computational model, Schlerf and colleagues (2013) reported that the slower learning rates in SCA are unlikely to be the result of increased motor noise. Indeed, when directly examined, minimal correlation is found between impairments in visuomotor learning and the severity of ataxia (Martin et al., 1996; Schlerf et al., 2013).

### Flexibility in aiming is not directly driven by sensory prediction errors

A second hypothesis is that sensory prediction errors are necessary to form an effective aiming solution. While some form of performance error would be needed to adjust aiming, it is unlikely that a sensory prediction error is the primary signal. A sensory prediction error occurs whenever there is a mismatch between the expected and actual consequences of a movement. This sensory prediction error is used to update a forward model to improve prediction. In a standard visuomotor rotation task, this error is defined as the difference between the cursor position and target (when taken as a proxy of the location for expected feedback). However, when participants use an instructed aiming strategy to counter the rotation, their hand slowly drifts away from the aiming location across trials (Mazzoni and Krakauer 2006). This occurs because a persistent sensory prediction error, the mismatch between the cursor position and aiming location in this context, continues to result in implicit updating of a forward model. Eventually, this implicit learning leads to poor performance (i.e., large target errors) and participants have to change their aim to offset continued implicit learning (Taylor and Ivry 2011). Models of performance in this aiming task suggest that changes in aim are driven by performance error, the difference between the target and cursor feedback locations, rather than sensory prediction error (Taylor and Ivry 2011; Taylor et al 2014). Sensory prediction errors, on the other hand, appear to lead to implicit learning even when irrelevant to the task goal (Mazzoni and Krakauer 2006; Schaefer and Thoroughman 2011; Morehead et al 2014).

Another problem with the hypothesis that the aiming deficit is related to an impairment in learning from sensory prediction errors is that the time course of aim reports looks quite different than that observed for implicit learning. First, with large perturbations (e.g., experiment 1), the aim reports are non-monotonic, with an initial large increase and then a gradual reduction over the course of the rotation block. Second, while the average data might look smooth, the aiming reports for individuals can change abruptly and in either direction, behavior that is reminiscent of exploration (see Taylor et al., 2014). This stands in contrast, to the slow, monotonic updating of an internal model that is considered to be the hallmark of motor adaptation (Huberdeau et al., 2015). Thus, it seems unlikely that the output of a forward model is the driving force for finding an aiming solution.

### An action-outcome maintenance account of impaired aiming in SCA

A third hypothesis considers less direct ways in which the cerebellum might support learning, and by extension, the cognitive capacity required for discovering an aiming solution. The cerebellum is known to be highly connected with much of the cerebral cortex, including prefrontal cortex (Buckner et al., 2011), and damage to the cerebellum can produce a constellation of neuropsychological impairments similar, albeit in milder form, to that observed in patients with lesions of prefrontal cortex (Buckner et al., 2013; Bodranghien et al., 2016). Drawing on ideas developed in perceptual domains (Cohen et al., 1997; Prabhakaran et al., 2000; Gazzaley et al., 2005), we have proposed that a network including prefrontal cortex and cerebellum forms something akin to a motor working memory system, one essential in the action domain (Ivry & Fiez, 2000; Spencer and Ivry, 2009). By this idea, the cerebellum helps represent and/or maintain task-relevant stimulus-response associations across trials. Theories of prefrontal cortex function have suggested a role in implementing and maintaining task sets across trials (Dosenbach et al., 2006 and 2007), possibly in a hierarchical manner where abstraction increases rostrally, allowing for the simultaneous search for action rules at multiple levels of abstraction (Badre and D’Esposito, 2009; Badre et al., 2010). By extension, the task set implemented by the prefrontal cortex would include the task relevant action-outcome associations, a network involving the cerebellum. Consistent with this theory, Spencer and Ivry (2009) showed that the impact of SCA on sequence learning was considerable when the task involved indirect, arbitrary S-R associations, but that SCA participants performed as well as controls in sequence learning when the S-R associations were direct.

An extension of this hypothesis may account for the aiming deficit observed in the current study. Converging on an appropriate aiming solution entails a cyclic process of hypothesis testing, generating possible solutions and then evaluating their efficacy. This process would require the maintenance of stimulus-response associations or, perhaps in the case of visuomotor rotations, action-outcome associations.

Moreover, the memory demands are likely greatly increased when there are eight target locations, especially with a visuomotor rotation (Krakauer et al., 2000) where implicit generalization appears quite narrow (Heuer and Hegele, 2011). Here an aiming hypothesis generated in response to an action at one target, may appear appropriate when applied to neighboring targets, but fail miserably when applied to distant targets. For example, aiming above the target location would be effective in countering a clockwise rotation for targets presented on the right side of the display but would be counterproductive if applied to targets on the left side of the display. Thus, the participant would need to maintain action-outcome associations at multiple target locations to eventually learn that the solution to the perturbation is common to all target locations in a polar coordinate reference frame. This idea would suggest that if the SCA participants had a compromised ability to maintain action-outcome association, then they would naturally have an impaired ability to counter a visuomotor rotation across the workspace.

Our action-outcome maintenance hypothesis makes three predictions. First, we would expect that the ability to employ an aiming solution might be related to the individual’s cognitive status. Second, we would expect that individuals with SCA would be able to develop an appropriate aiming solution if the perturbation was simpler. For example, these individuals might be able to use aiming to compensate for a translational perturbation. Third, we would predict that individuals with ataxia would disproportionately benefit if the demands on working memory were reduced, say by displaying the previous aim choice for each target, or by the use of only a single target location. These last two predictions can motivate future work on the multi-faceted contribution of the cerebellum to sensorimotor learning. At present, the results presented here underscore a role for the cerebellum, not only in implicit aspects of motor performance, but also when those tasks require a more explicit association to link a stimulus with an appropriate response to meet task goals.

Here we observed a dual-deficit in sensory-prediction-error-based learning (e.g., a forward model) and aiming. While at first glance the latter may seem odd, it perhaps shouldn’t be all that surprising given the cerebellum’s involvement in learning for many different types of tasks, from eye-blink conditioning to sequence learning. Indeed, the nearly uniform circuitry of the cerebellum, along with its connections to many areas of the rest of the brain (Buckner et al., 2011), suggests that it may be contributing to learning processes in a generalized manner that remains to be determined.

## Acknowledgements

We thank Sam McDougle for feedback on the manuscript. P.A.B and J.A.T. were supported by R01NS084948 from the National Institute of Neurological Disorders and Stroke and the Princeton Neuroscience Institute’s Innovation Fund. R.B.I. was supported by National Institutes of Health grants NS074917 and NS092079. The authors declare no competing financial interests.

